# Vector-clustering Multiple Sequence Alignment: Aligning into the twilight zone of protein sequence similarity with protein language models

**DOI:** 10.1101/2022.10.21.513099

**Authors:** Claire D. McWhite, Mona Singh

## Abstract

Multiple sequence alignment is a critical step in the study of protein sequence and function. Typically, multiple sequence alignment algorithms progressively align pairs of sequences and combine these alignments with the aid of a guide tree. These alignment algorithms use scoring systems based on substitution matrices to measure amino-acid similarities. While successful, standard methods struggle on sets of proteins with low sequence identity - the so-called twilight zone of protein alignment. For these difficult cases, another source of information is needed. Protein language models are a powerful new approach that leverage massive sequence datasets to produce high-dimensional contextual embeddings for each amino acid in a sequence. These embeddings have been shown to reflect physicochemical and higher-order structural and functional attributes of amino acids within proteins. Here, we present a novel approach to multiple sequence alignment, based on clustering and ordering amino acid contextual embeddings. Our method for aligning semantically consistent groups of proteins circumvents the need for many standard components of multiple sequence alignment algorithms, avoiding initial guide tree construction, intermediate pairwise alignments, gap penalties, and substitution matrices. The added information from contextual embeddings leads to higher accuracy alignments for structurally similar proteins with low amino-acid similarity. We anticipate that protein language models will become a fundamental component of the next generation of algorithms for generating MSAs.

Software availability: https://github.com/clairemcwhite/vcmsa

## INTRODUCTION

Multiple sequence alignments (MSAs) underlie much of sequence analysis in computational biology. Protein MSAs are the main input to algorithms for reconstructing gene and species trees (Felsenstein, 2004); for identifying conserved (Capra and Singh 2007), co-evolving (Marks et al. 2011), specificity-determining (Capra and Singh 2008) and positively selected sites (Miyata and Yasunaga 1980); for building protein motif and domain representations (Sonnhammer et al. 1998); and for predicting *de novo* protein structural features (Rost and Sander 1994) and full protein structure prediction (Jumper et al. 2021). Highly accurate MSAs are necessary, as misaligned amino acids can lead to incorrect inferences for this wide range of downstream tasks. However, when the average sequence identity amongst the sequences to be aligned is low, most MSA methods struggle to obtain high-quality alignments (Nute, Saleh, and Warnow 2019); sequence identity below 20-35% has been termed the “twilight zone” of alignment (Rost 1999).

Over the last 40 years, numerous algorithms for obtaining MSAs have been developed and refined, including CLUSTAL (Sievers et al. 2011), MAFFT (Katoh and Standley 2013), Muscle (Edgar 2004), T-coffee (Notredame, Higgins, and Heringa 2000), ProbCons (Do et al. 2005), Saté (Liu et al. 2012) and PASTA (Mirarab et al. 2015), among others. The predominant approach to align multiple sequences is to first align pairs of sequences, and then progressively combine these subalignments into larger alignments in an order determined by a guide tree (Feng and Doolittle 1987). Pairwise alignment algorithms (Smith and Waterman 1981; Needleman and Wunsch 1970) uncover a highest scoring alignment between two sequences when given a similarity measure between all pairs of amino acids and a gap penalty function.

Similarities between each pair of the 20 amino acids are typically based upon substitution matrices that encapsulate the frequency with which amino acid pairs are observed in equivalent positions across large datasets of alignments of homologous sequences (Altschul 1991; Dayhoff, Schwartz, and Orcutt 1978; Henikoff and Henikoff 1992). While in theory optimal pairwise alignment algorithms can be adapted to multiple sequences, the runtime would be exponential in the number of sequences, and further it is not clear how to score multiple amino acids in a single aligned column. Instead, guide-tree-based MSA approaches are heuristics that optimally combine pairs of alignments where the similarity of a pair of columns in two different alignments is computed based on substitution matrix scores of the amino acids within the columns. Alternatively, consistency-based progressive MSA approaches use similarity scores that consider how frequently pairs of amino acids in two different sequences are aligned to the same amino acid position in other sequences when considering optimal pairwise alignments (Notredame, Higgins, and Heringa 2000; Notredame, Holm, and Higgins 1998).

Here we consider a new approach to determining the similarities between amino acids across proteins, based upon protein language models (Chowdhury et al. 2021; Elnaggar et al. 2020; Rao et al. 2019; Bepler and Berger 2021; Rives et al. 2021). Protein language models are “self-supervised” deep learning language models that have been pre-trained on large compendia of protein sequences, generally trained to predict “masked” amino acids within input protein sequences based on the rest of the sequence. Once trained, these models embed each amino acid within a protein sequence as a high dimensional vector, and importantly these vectors capture the sequence “context” of each amino acid. Remarkably, machine learning models using these embeddings have been successfully trained to predict many protein structural and functional features, including conservation (Marquet et al. 2022), ligand binding (Littmann et al. 2021), homology (Heinzinger et al. 2022; Rives et al. 2021), and subcellular localization (Stärk et al. 2021). Since amino acids at equivalent positions in two different protein sequences play the same structural and functional role, we reasoned that clustering amino acid context vectors across sequences could be the basis of a new approach for aligning sequences. While recently neural network machinery has been leveraged to perform dynamic program based alignments (Morton et al. 2020; Petti et al. 2021), our approach is, to the best of our knowledge, the first that uses protein sequence embeddings to identify analogous positions across protein sequences in order to determine MSAs. Our MSA approach is based on clustering the amino acid contextual vectors produced by protein language models, and then using graph-theoretic approaches to determine a consistent ordering of MSA columns. Our method vcMSA (**v**ector-**c**lustering **M**ultiple **S**equence **A**lignment) is a true multiple sequence aligner that aligns multiple sequences at once instead of progressively integrating pairwise alignments. Our core methodology diverges from standard MSA methods in that it avoids substitution matrices and gap penalties, and in most cases does not utilize guide tree construction. In proof-of-concept testing on a set of sequences from 147 homologous protein families from the Quantest2 (Sievers and Higgins 2020b) dataset, we demonstrate that vcMSA tends to produce better alignments than multiple state-of-the-art alignment algorithms. Further, vcMSA excels when considering sequence families with highly divergent sequences. Thus, our protein language powered approach vcMSA obtains better alignments for those low identity sequence families that current methods have the most difficulty on.

## METHODS

### Overview

A brief overview of our algorithm is as follows. First, for all protein sequences, we obtain per-position and sequence-level embeddings via a protein language model (**Figure 1A**). Second, we cluster protein sequences via the sequence-level embeddings to uncover sets of sequence-similar proteins (**Figure 1B**). Third, within each cluster of sequences, we measure the similarities of amino acid vectors and for each amino acid find the one that is most similar to it in each of the other sequences (**Figure 1C**), and filter to amino acid pairings that are reciprocal best hits (RBHs) of each other. Fourth, we build a RBH network and cluster the network to find “guidepost” amino acid positions in different sequences that are clearly aligned to each other (**Figure 1D**). Fifth, we order the clusters into columns of the MSA by constructing a directed acyclic graph based on sequential positions within each protein sequence and performing a topological sort (**Figure 1E**). Sixth, we use guidepost clusters to limit the scope of searches for remaining unplaced amino acids (**Figure 1F**). Seventh, we continue creating guideposts and assigning amino acids to columns until all amino acids are placed in the alignment (**Figure 1G**). Finally, we combine sub-alignments from the sequence clusters into the final MSA (**Figure 1H**). Further details about each of these steps is provided below.

**Figure 1.**
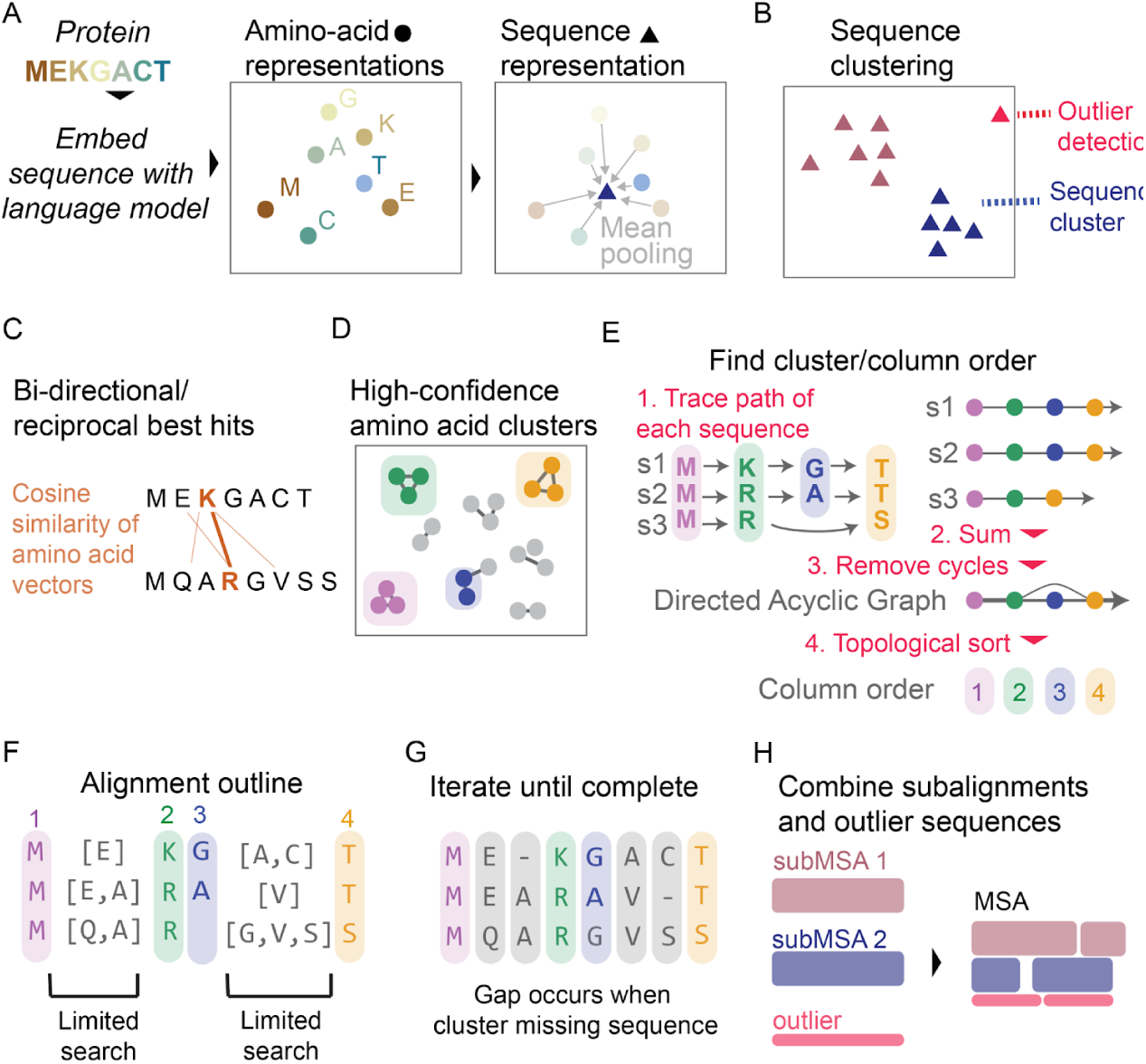
Overview of vcMSA algorithm. (A) Proteins are embedded using a protein language model to produce vector representations of each amino acid, and the mean of these amino acid embeddings is taken to produce a sequence-level representation. (B) We cluster sequence representations, and detect outlier sequences. (C) For each sequence cluster, we determine bi-directional/reciprocal best hits of cosine-similarity between pairs of amino acids in different sequences. (D) From a network built from reciprocal best hits, we determine confident clusters of amino acids, corresponding to columns in the MSA. (E) To determine column order, we trace the path of each sequence through clusters and combine all paths into one network, taking edge weights from the number of sequences which traverse between the pairs of clusters. We trim any clusters which cause cycles, and use a topological sort of the resulting directed acyclic graph to find column order. (F) Clusters/columns limit scope of search for unplaced amino acids. (G) We iterate limited searches until all amino acids are placed. Gaps in the alignment occur when a cluster does not contain an amino acid from a sequence. (H) We combine alignments from each sequence cluster and outliers in the final output MSA.

### Embedding generation

For each protein sequence, we use the encoder portion of the ProtT5-XL-UniRef50 language model to generate sequence embeddings (Elnaggar et al. 2020). T5 models undergo self-supervised training on a large corpus of text, in this case over 11 million protein sequences in the Uniref50 data set (Elnaggar et al. 2020). For each amino acid in a sequence, the ProtT5-XL-UniRef50 model produces one embedding vector of length 1024 for each of its 24 encoder layers. We choose to use embeddings from the final 16 layers, giving us a final vector representation of each amino acid with dimension 16384. For each sequence, we average all of the vector embeddings for each of its amino acids to obtain a sequence-level embedding. We additionally add padding to sequences, where we first add ten X’s to the start and end of each sequence prior to embedding (**Supplemental Figure 1A**).

### Sequence clustering

We find that vcMSA performs best on semantically consistent sets of sequences, and that performance is degraded when outlier sequences are included. Therefore, we first cluster input sequences based on sequence-level embeddings to obtain groups of similar sequences, and detect outliers (**Supplemental Methods**). We then apply the remaining steps of vcMSA to each cluster of sequences individually. Left out sequences and alignments for clusters are merged into a full alignment as described below.

### Amino acid similarity across sequences

The contextual similarity between a pair of amino acids is defined as the cosine similarity between their embedding vectors. For each amino acid in each sequence, we find the most similar amino acid in each of the other sequences. We use Facebook’s fast AI similarity search (faiss) library to perform nearest neighbor searches for all amino acids (J. Johnson, Douze, and Jégou 2017). For each pair of sequences, the set of pairwise similarities between amino acids between different sequences is filtered to only RBHs, where the best match to a query amino acid in the target sequence must also have its highest score in the query sequence be the query amino acid (**Figure 1C**). For each pair of sequences, we consider all RBHs between amino acids, and keep a largest set of these that are “consistent” with each other in a pairwise alignment using a longest increasing subsequence formulation. In particular, if the *i*-th and *j*-th amino acids in the first sequence (we assume without loss of generality that *i* < *j)* have as their RBHs the *k*-th and *l*-th amino acids respectively in the second sequence, then this pair of RBHs is consistent only if *k* < *l*; that is, it is possible for a pairwise alignment between these two sequences to simultaneously align the *i*-th and *j*-th amino acids in the first sequence with the *k*-th and *l*-th amino acids in the second sequence. For each amino acid *i* in the first sequence, we label it with *s_i_*, the position within the second sequence of its RBH, and then uncover the longest increasing subsequence of the *s_i_*; the RBHs corresponding to the *s_i_* that are part of the longest increasing subsequence are all consistent with each other. The longest increasing subsequence can be found in 𝑂(𝑛 𝑙𝑜𝑔 𝑛) where 𝑛 is the length of the sequence (Fredman 1975). We remove from consideration any RBHs that are not part of the longest increasing subsequence; in this manner, we obtain a filtered set of RBHs across all pairs of sequences.

### Amino acid clustering

We build a network where there is a node for each amino acid in each sequence, and there is an edge between two nodes if the corresponding amino acids are RBHs included in the filtered set of RBHs. We use this network of RBHs in order to cluster sets of amino acids that align to each other with high confidence (**Figure 1D**) (**Supplemental methods**). Amino acids within a cluster will comprise a column in the MSA. We term these aligned positions guideposts, and will use these guidepost columns to limit the scope of future searches for amino acids that align to each other (**Figure 1F**). Our guidepost columns are similar to user-defined or automatically inferred “anchor points” that have been utilized in the past to constrain traditional MSA approaches (Morgenstern et al. 2006; Pitschi, Devauchelle, and Corel 2010).

### Column order determination

Each cluster corresponds to a column of the multiple sequence alignment; however, these columns must be placed in the correct order in the alignment that is being built. To do this, we will first build a graph based on the correct ordering of amino acids in each protein sequence, and then prune this graph to be a directed acylic graph (DAG) and perform a topological sort of this graph (**Figure 1E**). In particular, we will introduce a node for each cluster. Next, for clusters 𝐴 and 𝐵, if there exists amino acids 𝑢 ∈ 𝐴 and 𝑣 ∈ 𝐵 where 𝑢 and 𝑣 are in the same sequence, 𝑢 is before 𝑣 in the sequence, and 𝑣 is the first clustered amino acid after 𝑢 in the sequence, then we will add a directed edge from 𝑢 to 𝑣. This edge will be weighted by the number of sequences where such 𝑢 and 𝑣 exist. Out-of-order amino acids, where an amino acid is placed in a cluster that occurs too late or too early in the MSA, will cause cycles in this directed graph. To remove these out-of-order amino acids, we detect feedback arc sets (Eades, Lin, and Smyth 1993; Csardi and Nepusz 2006), finding edges to remove to obtain an acyclic graph and prioritizing edges with low weight for removal. Since feedback arc set detection is NP-complete, we use the linear time heuristic approach of Eades, Lin, and Smyth 1993. Feedback arc set detection has been previously used in MSA to enforce order consistency for multiple local areas of alignment (Pitschi, Devauchelle, and Corel 2010).

For each node-pair of each feedback arc, we discard the node/cluster ID with lowest total edge weight, and finally confirm that the trimmed network is a directed acyclic graph. We then perform a topological sort of the graph to obtain an ordered path that visits every node (cluster ID). This sequence of cluster IDs is the order of columns in the multiple sequence alignment.

### Limited scope searches

At this stage, we have a subset of guidepost columns in the alignment, and sets of amino acids which have not been assigned to any column (**Figure 1F**). If an unassigned amino acids occurs after an amino acid which has been assigned to column 𝑝, and before an amino acid which has been assigned to column 𝑞, we only need to consider amino acids from other sequence that also fall between 𝑝 and 𝑞 as candidate matches.

We repeatedly alternate two phases of adding amino acids into the alignment until there are no remaining unplaced amino acids with cosine similarity > 0.1 to any potential match amino acid in the scope. Our two phases are 1) create new columns by amino acid clustering, as described above, among all amino acids also falling between the appropriate guideposts and 2) add amino acids into existing columns by best match score (**Supplemental Methods**). Existing columns can only be modified during the second phase with the addition of new members. In early iterations following clustering stages, we remove clusters where any member of the cluster is “stranded”, meaning that neither of its previous or next amino acids is found in the previous and next aligned column/cluster. This quality control step prevents early errors where an amino acid is placed in the wrong cluster. For the final stage, amino acids with no matches to any clusters or other amino acids in the scope are placed in their own cluster of size 1.

### Gaps

Our method does not require parameters related to setting gaps. Once all amino acids are assigned to columns, we lay out the final alignment (**Figure 1G**). A column that has one amino acid from every sequence has no gaps. If a column does not contain an amino acid from a particular sequence, a gap is left for that sequence in that column. Our algorithm therefore does not need to use gap opening or gap extension parameters which are required by other methods.

### Overclustering

We observe cases of overclustering, where two adjacent columns contain amino acids from a mutually exclusive set of sequences. We detect and merge these paired columns using a low threshold cosine similarity score of 0.1 between amino acids in adjacent clusters, on the basis that independent insertions of one amino acid at the site is biologically unlikely.

### Benchmarking

We use the Quantest2 (Sievers and Higgins 2020) reference alignment dataset, which consists of 151 protein sets, each composed of 1000 proteins. The Quantest2 alignment set is a subset of the HOMSTRAD (Stebbings and Mizuguchi 2004) database of curated structure alignments. We use HOMSTRAD alignments that are not included in the Quantest2 set to evaluate correspondence between Reciprocal Best Hit pairs of amino acids and aligned positions. We modified the nextflow pipeline `nf-benchmark` (Garriga et al. 2019) to wrap our scripts and manage comparison to other methods and evaluation on the Quantest2 set of alignments.

We compare our method to Clustal Omega (Sievers et al. 2011), MAFFT-FFTNS1, MAFFT-GINSI, MAFFT-LINSI (Katoh and Standley 2013), MUSCLE (Edgar 2004), ProbCons (Do et al. 2005), T-Coffee (Notredame, Higgins, and Heringa 2000), UPP (Nguyen et al. 2015), and FAMSA (Deorowicz, Debudaj-Grabysz, and Gudyś 2016), and build initial guide trees for all methods except UPP with MAFFT-PartTree (Katoh and Toh 2007).

We use the same default parameters for other algorithms as in the Supplementary Information of Garriga et al. 2019. As with previous work, alignment performance is evaluated using gold-standard reference alignments of the first three proteins in each fasta file (Garriga et al. 2019). For the proof-of-concept testing described here, we created smaller sets of proteins to align consisting of 20 sequences, where we select the first three sequences from each fasta file, as well as 17 randomly selected protein sequences. We additionally remove four protein sets with over four-fold difference in length between the three reference sequences and the 17 random sequences, leaving a benchmarking dataset of 147 protein sets. We calculate sequence identity for each gold standard alignment by recording the mean proportion of times each aligned position in the pairs of sequences match exactly. For sequence identity bins, identities <0.2 fall in bin 0.1, <0.3 fall in bin 0.2, and so on. For runtime measurements, we also create larger sets of proteins consisting of 50 and 100 sequences, where we select the first three sequences and then 47 and 97 sequences, respectively.

For evaluating against structured portions of the alignment, we filter down gold standard alignments to the subset of amino acids with DSSP structure codes H, G, I, E, and B (Kabsch and Sander 1983).

### Evaluation score

We use the Total Column score calculated with the aln_compare plugin from T-Coffee to score sequence alignments against the gold standard alignments. This score is the percent of columns in the output alignment which fully match columns in the reference gold standard alignment.

### Speed

On a NVIDIA A100 GPU, initial protein embedding of 20 sequences takes on average 22.6 seconds, and 100 sequences takes on average 31.3 seconds, including loading the model. On a CPU configuration, protein embedding is substantially slower, taking two to five minutes to embed 20 sequences, depending on the length of the sequences and CPU memory.

The time to produce an alignment depends on the number of iterations required to place all amino acids. An alignment of 20 sequences on a CPU with 32 GB of memory takes between 15 seconds and 5 minutes, for 50 sequences between 1 minute and 10 minutes, and for 100 sequences between 5 and 16 minutes (**Supplemental Figure 2**).

As our implementation of the vcMSA is a proof-of-concept, we expect the speed of producing alignments to fall substantially with program optimization.

### Merging subalignments and excluded sequences

13/147 of our Quantest2 20 protein sets contain more than one cluster of sequences and 59/147 contain at least one outlier sequence.

For these 63 protein sets, we must combine vsMSA produced subalignments and excluded sequences into a complete alignment containing all sequences (**Figure 1H**). For this step, we use MAFFT-LINSI merge (Katoh and Standley 2013), which uses our sets of pre-aligned sequences as internal nodes of a guide tree constructed by UPGMA, using the command `mafft --clustalout --merge key_table --auto sub.alns > merged.aln`. For the remaining 84 protein sets, no merging of alignments is necessary. While in a typical use case, outlier sequences would be removed, we include them for compatibility with benchmark protein families.

### Code availability

We provide a python package ‘vcms’ at the https://github.com/clairemcwhite/vcmsa Github repository. Our modified version of the `nf-benchmark` (Garriga et al. 2019) is available at https://github.com/clairemcwhite/nf-benchmark-vcmsa

## RESULTS

### Reciprocal best hits between amino acid representations are a solid foundation for multiple sequence alignment

To demonstrate that reciprocal best hits between amino acid embeddings in two different sequences accurately reflects aligned amino acids, we measured the proportion of aligned positions in 562 pairs of sequences from the HOMSTRAD gold standard structural alignment database (Stebbings and Mizuguchi 2004) that are reciprocal best hits when comparing amino acid embeddings. For 93.2% of gold standard protein alignments, at least 50% of aligned columns are additionally reciprocal best hits between amino acid embeddings (**Figure 2A**). The correspondence between reciprocal best hits and aligned columns is roughly dependent on sequence identity of the alignment (**Figure 2B**). Thus, protein language models are highly effective in identifying analogous positions across protein sequences. These reciprocal best hits are the main input to the vcMSA algorithm.

**Figure 2.**
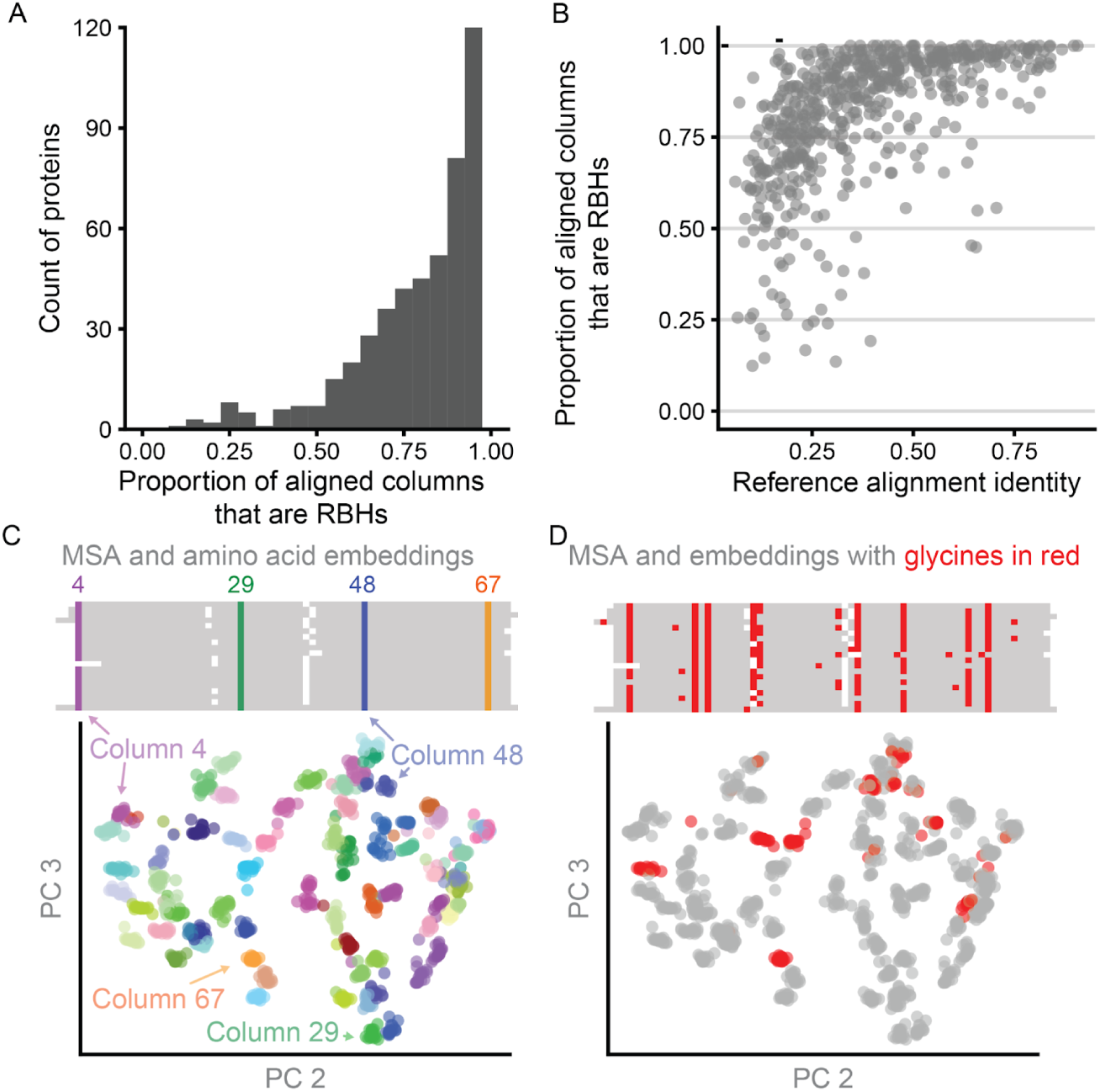
Reciprocal best hits and clustered amino acid representations reflect aligned columns. (A) For 562 pairs of aligned proteins from the HOMSTRAD database, reciprocal best hits between vector representations frequently correspond to correctly aligned positions. For each gold standard alignment, we compute the fraction of aligned gold standard positions that are additionally reciprocal best hits for that protein pair, and display these results as a histogram. (B) The proportion of columns in the gold standard pairwise alignments that are reciprocal best hits is related to sequence identity of the reference alignment. (C) Principal components 2 and 3 for amino acid vector representations of twenty cold shock proteins from the csp protein family. Amino acid representations (below) are colored by their corresponding column in the multiple sequence alignment (above). (D) Identical amino acids in different columns are distinguishable from each other. All glycines in the amino acid PCA plot and the multiple sequence alignment are colored red.

### Clusters of amino acid representations correspond to columns in the Multiple Sequence Alignment

When amino acid representations in a PCA are colored by their corresponding column in the multiple sequence alignment, it can be seen that amino acids in the same aligned column cluster together (**Figure 2C**). We show this correspondence with twenty sequences from the csp protein family using their HOMSTRAD alignment. One initial concern about using amino acid representations for multiple sequence alignment was that the representations of the same amino acid (e.g., glycine) would be too similar wherever that amino acid occurs in the sequence. Instead we find that representations of the same amino acids are distinguishable across the alignment (**Figure 2D**).

### vcMSA produces more accurate alignments than other algorithms, particularly at low sequence identity

We benchmarked vcMSA on 147 protein alignments consisting of 20 sequences each from the Quantest2 dataset, and compared our results to seven alignment algorithms (**Figure 3A**). As the metric to evaluate alignment accuracy, we chose Total Column score, which measures the number of columns that match the gold standard alignment, due to its high dynamic range relative to other metrics. vcMSA generally matches or exceeds the alignment accuracy of other algorithms, and performs especially well on the lowest sequence similarity proteins in the benchmark. As one striking example, vcMSA is able to align the Asp_Glu_race_D family with a Total Column score of 61.7, while the maximum score of other methods is 0.9.

**Figure 3.**
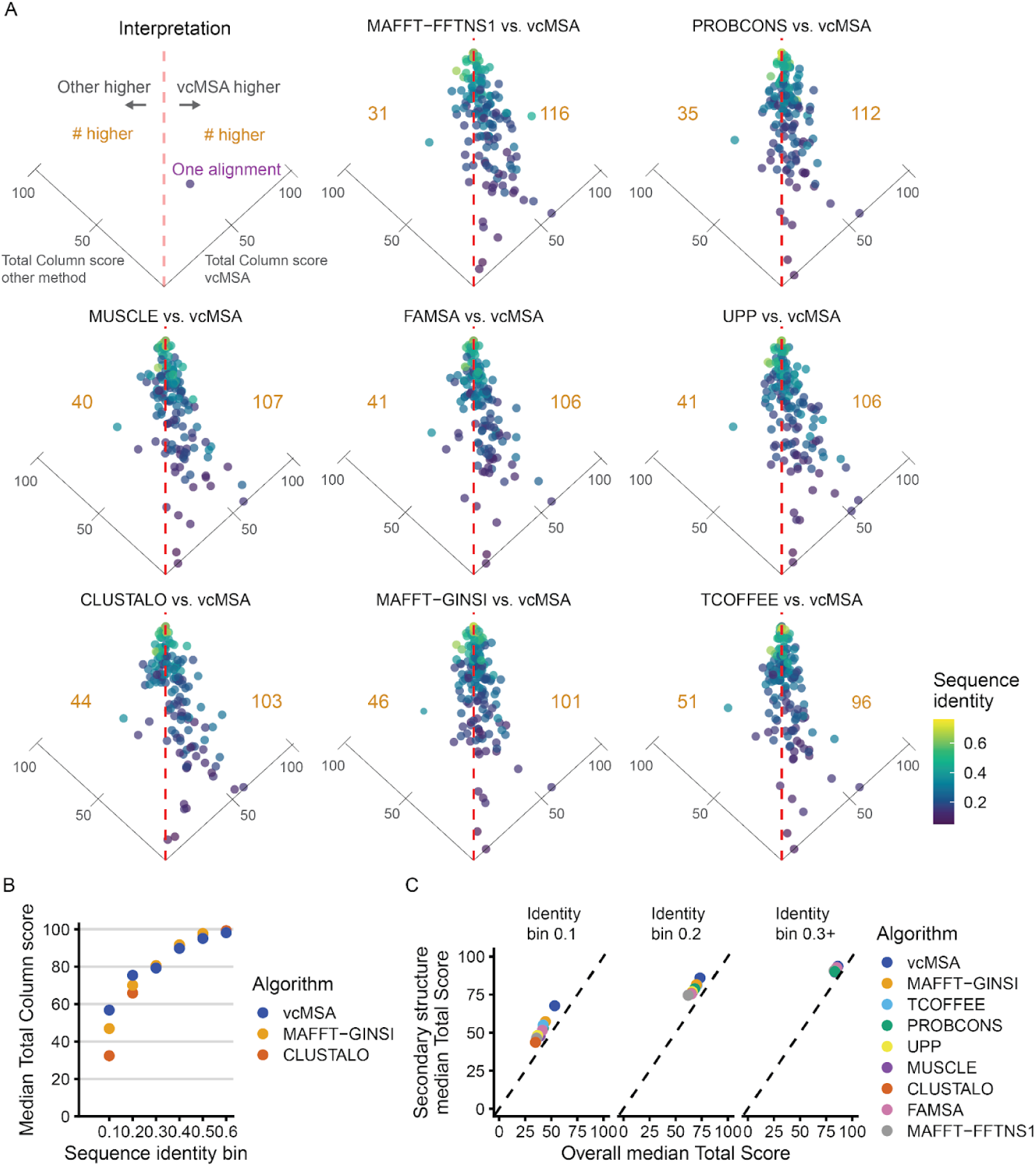
vcMSA alignments are frequently more accurate than previous methods, particularly for alignments with low levels of sequence identity. We benchmarked our implementation of vcMSA on a reference set of 147 gold standard multiple sequence alignments from the Quantest2 dataset. (A) We compare our method to eight state-of-the-art algorithms for sequence alignment. Each point corresponds to one protein alignment, and is colored by its sequence identity. Each panel is a tilted scatterplot of relative performance of each other algorithm tested against vcMSA by Total Column score of produced alignments. Points to the right of the vertical dashed line are alignments where vcMSA outperformed the other method. The number of alignments to the left or right of the line is in gold. vcMSA performs better on more test sets than all other tested algorithms. The point lying on the right axis line is the Asp_Glu_race_D alignment, which all other algorithms perform poorly on (TC score ∼0). (B) We compare the median Total Column score of vcMSA alignments to MAFFT-GINSI and Clustal Omega alignments, grouping alignments by sequence identity bins. We observe increased scores at low sequence identities. (C) We show median Total Column scores for structured portions and full gold standard reference alignments for all algorithms, separated by sequence identity bin.

For low sequence identity alignments, we frequently exceed the alignment accuracy of existing algorithms. In our benchmarking, as compared to other methods, we perform notably better on low sequence identity alignments. **Figure 3B** illustrates the performance of vcMSA against the two algorithms which score highest and lowest on sequences with identity < 0.2, MAFFT-GINSI and Clustal Omega, respectively. As sequence identity increases, vcMSA generally matches the accuracy of other algorithms. When we compute Total Column score for structured portions of the gold standard alignment, compared to the full gold standard alignment, it is apparent that vcMSA performs better on structured components (**Figure 3C**).

Further, vcMSA is specifically strong at structured regions of the lowest identity protein sets. This is likely due to the language model capturing structural properties of these low identity structured regions, even though their sequence identity is beyond the typical twilight zone limit.

## DISCUSSION

While sequence aligners were some of the earliest bioinformatic algorithms, and sequence alignment forms the basis of many computational biology analyses, multiple sequence alignment is not a solved problem. Current sequence alignment algorithms particularly struggle with alignments in the so-called twilight zone of sequence identity, where structure and function is conserved, but amino acid sequence is not.

Here, we present vcMSA, a novel algorithm for multiple sequence alignment that diverges substantially from other approaches, and demonstrate that it is an improvement on the state-of-the-art for some of the most challenging to align protein families. The ability to create accurate multiple sequence alignments opens up these low sequence identity families to all the downstream bioinformatics applications that take alignments as input. Notably, the core vcMSA algorithm avoids standard features of the most widely used alignment algorithms, including gap penalties, substitution matrices, and guide trees. While substitution matrices and gap penalties are standard components of alignment approaches, they are used to formalize optimization criteria and do not reflect fundamental properties of alignments. Instead, aligned amino acids within an MSA are meant to be those that have evolved from a shared ancestral amino acid.

Protein language models may be especially suited to identify these analogous amino acids, as they capture structural and functional properties of amino acids via sequence context, much as natural language transformer models capture the semantics of words in a sentence based on other words (Vaswani et al. 2017). Interestingly, we find that vcMSA performs particularly well at aligning structured portions of sequences; it is possible that the less well structured portions of sequences do not have specific functionally analogous positions in other sequences.

Protein alignment based on protein language representation appears to be most suited to global alignment for sets of protein sequences with similar structure and function, where the underlying meaning of the sequence is as consistent as possible. We have observed that alignment of a small subsequence of a protein (e.g., one domain) to a longer protein is often highly inaccurate, as the context embeddings for the small portion are sufficiently different from those computed for the longer protein. To produce the highest quality alignment, outlier sequences can be automatically detected based on sequence embeddings, and removed from the alignment process. In these cases, we first use sequence representations to divide sequences into high similarity groups, and detect outlier sequences. We then perform vcMSA on each cluster of sequences individually, producing high quality sub-alignments. While we currently use MAFFT-LINSI to combine sub-alignments produced by vcMSA, we are exploring methods to combine sub-alignments using purely language models. However, divergence in protein meaning between sequence groups complicates this process.

Additionally, for alignments where start positions are not aligned, we find it helpful to add sequence padding to either end of the sequence prior to embedding, which reduces an observed first character effect, where the first characters of each sequence tend to have the most similar embeddings, regardless of the sequence context of that first character (**Supplemental Figure 1A**). Interestingly, in these cases, padding can substantially improve correctly aligned columns beyond the first column (**Supplemental Figure 1B**). However, adding padding either does not change or substantially degrade alignment accuracy for protein sets where the start positions align.

Interestingly, for certain protein groups, we observe a sequence-level offset in amino acid embeddings. Batch correction by sequence performed with ComBat (W. E. Johnson, Li, and Rabinovic 2007) can reduce these batch effects (**Supplemental Figure 1C**). When present, these batch effects are most damaging for small sets of sequences. For example, three sequences of Glyco_hydro_18_D2 which show sequence-level batch effects cannot be accurately aligned without batch correction. However, for larger numbers of sequences, batch correction has a negligible effect on alignment accuracy (**Supplemental Figure 1D**). Further research is needed to understand where these sequence-level batch effects originate from, and how to recognize and reduce them when comparing amino acid embeddings across sequences.

As our implementation is a prototype, we have not substantially optimized our scripts for speed and memory usage, with much room for improvement in these areas. To identify similar amino acid positions at-scale between larger numbers of sequences, we can use tools developed for ultra fast nearest neighbor searches of millions and billions of vectors (Johnson, Douze, and Jégou 2017). Additionally, our algorithm’s runtime is greatly dependent on the number of iterations required to place all amino acids, which is related to the number of clustering errors (when an amino acid is sorted into a cluster that conflicts with the order of amino acids in the sequence). Currently, we use the very basic approach of finding connected components and choosing them to be clusters; we believe that slightly more sophisticated approaches (e.g., those requiring a higher edge density within clusters) would reduce errors in clustering and dramatically decrease runtimes.

There are also numerous ways in which the algorithm can be improved to obtain better alignments. For example, while here we use reciprocal best hits to identify amino acids that align to each other, and disregard similarities to other amino acids in the sequences, an approach that additionally considers near-best hits along the sequence is likely to perform better; in this case, the weighted version of maximal noncrossing matching (Malucelli, Ottmann, and Pretolani 1993) could find multiple analogous positions at once in a manner that respects positional ordering and could also remove spurious similarities between pairs of amino acids.

Our prototype implementation of the vcMSA algorithm outperforms existing alignment algorithms at aligning a benchmark of sets of proteins. It is particularly successful at aligning groups of low similarity sequences, filling a gap in the capabilities of other algorithms. Another advantage of using amino acid embeddings is that we can additionally score the confidence of each column in the alignment, by measuring average cosine similarity of amino acid embeddings from the same column. This confidence score will allow filtering of alignments to only confidently aligned positions in downstream tasks such as phylogenetic tree building.

Here we demonstrate both the utility of protein language embeddings in multiple sequence alignment and a new multiple sequence alignment algorithm fully orthogonal to existing approaches. We expect accuracy to only improve with optimized choice of amino acid representations and the development of larger language models which will more richly capture the identity of each amino acid. Overall, we anticipate that protein language models will play an important role in the next generation of methods for determining multiple sequence alignments.

## ACKNOWLEDGMENTS

CDM acknowledges support from the Lewis-Sigler Institute for Integrative Genomics. MS acknowledges support from the NIH. The authors thank Joshua Akey for helpful discussions, and also thank Princeton University High Performance Computing Center for providing high-performance computing resources that have contributed to the research results reported in this paper.

## SUPPLEMENTAL FIGURES

**Supplemental Figure 1:**
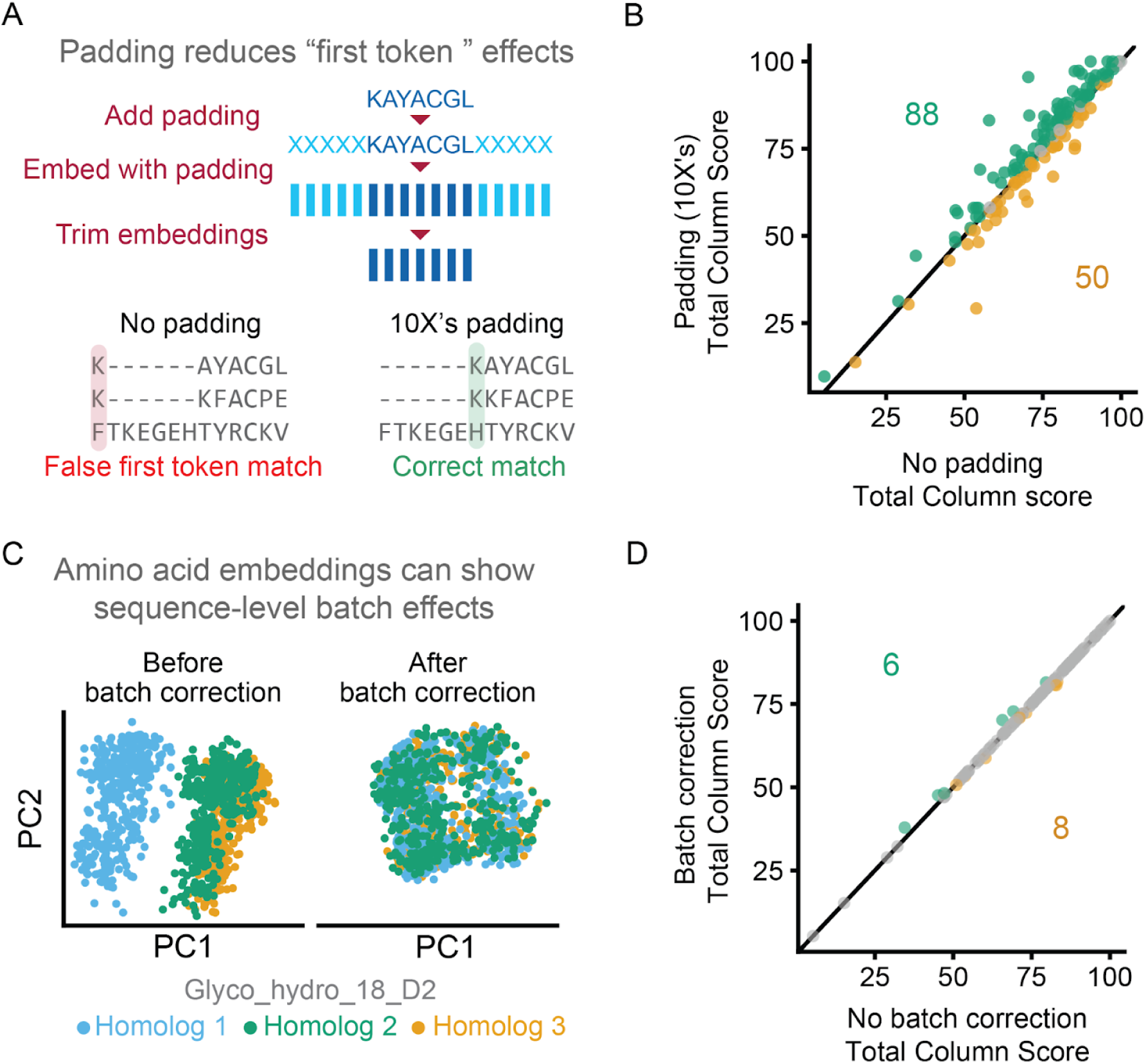
Illustration of sequence padding and batch correction. (A) For sequence padding, ten X’s are added to the start and end of each sequence prior to embedding. Embedding positions corresponding to these twenty X’s are trimmed prior to use. (B) Adding padding generally increases the Total Column scores for the benchmarking set of 147 alignments of 20 proteins. (C) Amino acid embeddings can show sequence-level batch effects. (D) For alignments of 20 sequences, batch correction has a minimal effect on alignment accuracy.

**Supplemental Figure 2:**
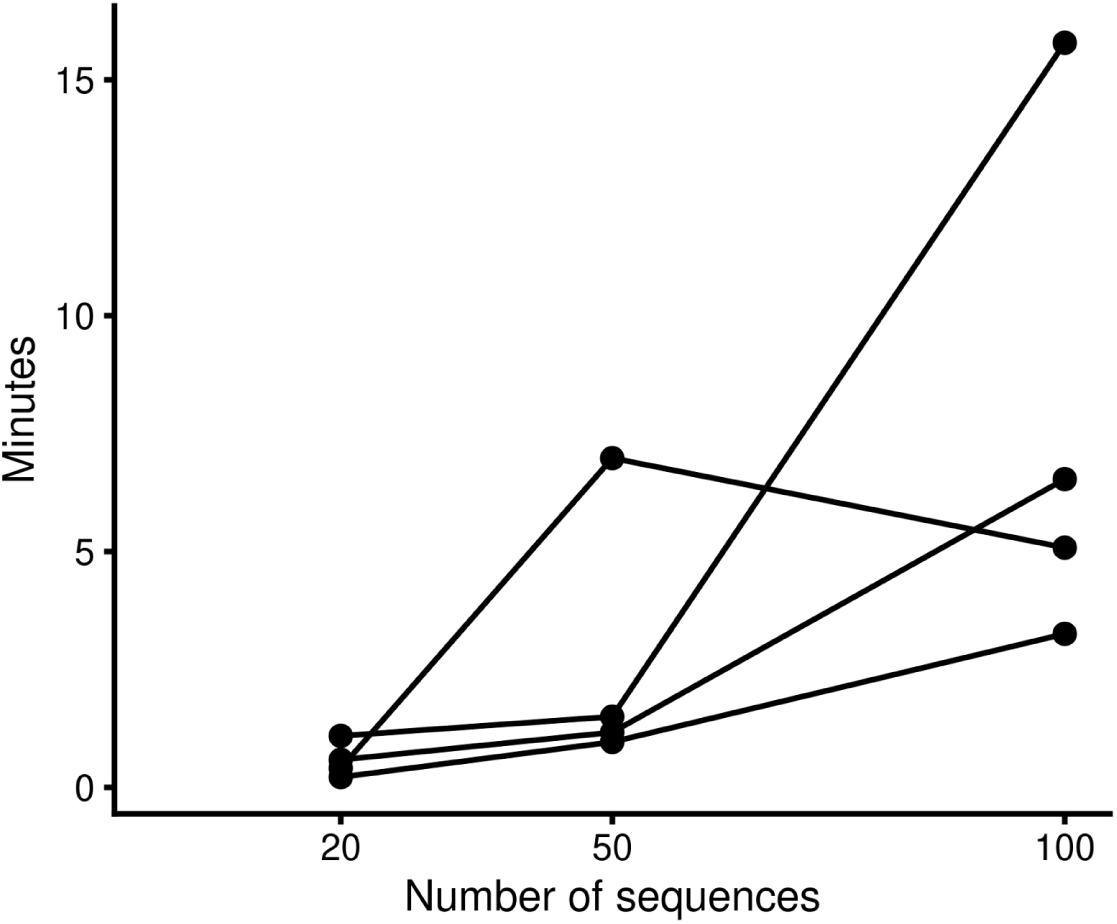
Relationship between number of sequences and time to complete an alignment. Each line represents one protein with 20, 50 and 100 sequences.

## SUPPLEMENTAL METHODS

### Sequence clustering and outlier detection

We build a network where there is a node for each sequence and edge weights between nodes correspond to the cosine similarities of their sequence-level embeddings and then cluster this network with the 𝐺. 𝑐𝑜𝑚𝑚𝑢𝑛𝑖𝑡𝑦_𝑤𝑎𝑙𝑘𝑡𝑟𝑎𝑝() function from the python-igraph package, which finds densely connected neighborhoods using random walks (Csardi and Nepusz, 2006). To remove outlier sequences, we sum the total cosine similarity of each sequence to the other sequences in its cluster. If a sequence has a z-score of less than -5 relative to the total cosine similarity of each of the other sequences, we remove it from the cluster and exclude it from the vcMSA alignment process.

### Amino acid clustering

To cluster sets of amino acids, we first simply extract individual connected components of the RBH network. If a connected component contains at most one amino acid from each sequence, and contains amino acids from over half the sequences, we align these amino acids to each other. In subsequent iterations of vcMSA, if a connected component contains more than one amino acid from the same sequence, we either remove the more weakly connected amino acid, or for larger connected sets, apply walktrap clustering to break it into smaller clusters. In early iterations, we discard small natural clusters, however reduce this minimum cluster size requirement over subsequent iterations, until clusters of size two are permitted. Finally, amino acids which are not placed in any cluster are placed in clusters of size one.

### Adding amino acids to existing clusters

During the second phase of the limited scope search, we add unsorted amino acids to existing clusters/columns. For example, in Figure 1F, the limited scope of the unsorted G, V, and S in sequence 3 includes the open position in cluster 3.

If an unsorted amino acid has a mean cosine similarity to an in-scope cluster’s amino acids greater than a threshold T (set to 0.5 in our experiments), that amino acid is added to the cluster.

### Batch correction

Batch correction of amino acid embeddings by sequence performed with ComBat (W. E. Johnson, Li, and Rabinovic 2007). Here, sequences are analogous to a batch, and amino acid embeddings are analogous to samples within that batch. Briefly, ComBat removes variations between batches using an empirical Bayes framework.

## Notes

### Competing Interest Statement

The authors have declared no competing interest.

### Summary of Updates

Updated text, additional supplemental figures

